# Cross-species multi-omics integration reveals lung alterations in acute kidney injury across mouse, pig, and human airway organoid models

**DOI:** 10.64898/2026.10.13.758311

**Authors:** Zhi Tian, Nohely Hernandez Soto, Diane Allen-Gipson, Georgina Agyekum, Xiaoqi Sun, Nadezhda N. Zheleznova, David Raj Chellappan, Nathan Hall, Kristof Williams, Runze Ni, Yuqian Tian, Sebastian Cuadrao, Vijay Subramanian, Lucian Lozonschi, Kiran Dhanireddy, Isabella Lozonschi, Carlos Esquivel, Colby L. Parris, Minh H. Tran, Lei Wang, Ruisheng Liu

## Abstract

**Background:** Acute kidney injury (AKI) is frequently associated with the development of distant acute lung injury (ALI); however, its broader clinical impact and the underlying temporal molecular mechanisms remain incompletely characterized.

**Methods:** We used aggregate deidentified Epic Cosmos data to describe all-cause mortality and hospital length of stay (LOS) in an AKI cohort (13,815,092 patients; 2016–2025) and a subgroup with subsequent lung injury (AKI-lung). Mechanistically, we integrated matched RNA-seq and proteomics from a hyper-acute pig model (4 h reperfusion), an established mouse model (24 h reperfusion), and primary lung organoids exposed to mtDAMP-conditioned media, using MOFA+, PROGENy, decoupleR/CollecTRI, and DIABLO. Candidates were validated in primary human donor lungs obtained from LifeLink, separated into Control (< 1.3 mg/dL, n = 5) and AKI (≥ 1.3 mg/dL, n = 5) groups based on the highest blood creatinine level measured in hospital.

**Results:** The AKI-lung subgroup had higher observed mortality than the overall AKI cohort (36.67% vs. 25.45%, p < 0.001). Mean hospital LOS was 11.3 versus 6.9 days in the reported AKI-lung and AKI alone queries, respectively (unadjusted p < 0.001). Multi-omics analyses cross-species indicated a temporal shift in proteogenomic coupling. In the hyper-acute pig model, we observed a largely post-translational response, including ciliary breakdown and metabolic changes. The established mouse model showed a coordinated transcriptional inflammatory response (r = 0.316), characterized by PI3K/Akt pathway activation and downregulation of vascular and immune regulators (Aplnr, Cd74). Human lung organoids exhibited transcriptional downregulation of tissue remodeling genes and persistent ATF6/ATF4-associated ER stress, consistent with a reparative phase under proteostatic strain. Unsupervised MOFA+ revealed conserved cross-species axes (Factors 6 and 7) linked to vascular repair and hemostasis. Upstream regulon analysis identified the endothelial integrity regulators, including FOXF1. Most importantly, targeted validation in primary human donor lungs confirmed a statistically significant, selective down-regulation of the core circadian regulator circadian locomotor output cycles kaput (CLOCK) transcript (p < 0.05).

**Conclusions:** Aggregate clinical findings suggest an association between lung injury following AKI and worse outcome. Kidney-Lung crosstalk initiates as hyper-acute post-translational microvascular breakdown in swine and rapidly transitions into localized uncoupling of cell-autonomous pulmonary circadian machinery (CLOCK) in clinical human AKI, offering human-validated targets to intercept distant organ failure.

## INTRODUCTION

Acute kidney injury (AKI) is a major global health concern, affecting over 13 million individuals worldwide each year and associated with considerable morbidity, mortality, and risk of progression to chronic kidney disease and other long-term complications ^1^. Once regarded as a primarily renal disorder, increasing clinical and experimental evidence now supports its association with a broad spectrum of distant organ injuries, including the lungs, heart, liver, and intestines ^2–6^. Of these, the lung appears particularly vulnerable, with AKI-associated lung injury linked to increased morbidity and mortality^7^. Despite the clinical importance of AKI-associated lung injury, the molecular mechanisms initiating kidney-lung communication and the temporal dynamics of this inter-organ crosstalk remain incompletely defined.

Recent studies suggest that mitochondrial damage-associated molecular patterns (mtDAMPs), released from injured kidneys, may act as endogenous signals that contribute to inflammatory responses in distant organs^3,8–10^. In the lung, these mediators are proposed to affect the alveolar-capillary barrier, activating inflammatory and stress pathways that may impair endothelial integrity, increase vascular permeability, and promote alveolar edema^11^. Such early events are associated with changes in endothelial signaling, vascular regulation, and cytoskeletal remodeling that can compromise barrier function ^12^. However, the initial molecular programs preceding immune cell infiltration remain insufficiently characterized. Capturing these transient responses is challenging due to their rapid onset and the limitations of conventional single-species models.

Addressing these early molecular events requires experimental systems that can capture both temporal dynamics and cross-species conservation of kidney-to-lung communication after AKI. While rodent models have provided critical insights into transcriptional and immunologic mechanisms, they may not fully reflect the hyperacute physiological and molecular responses seen in large animals, whose cardiovascular and pulmonary physiology more closely approximate that of humans ^13–15^. To improve relevance to human disease, complementary platforms using human-derived tissues and organoids can help interrogate epithelial stress responses with reduced confounding from systemic inflammation^16–18^. Therefore, integrating temporally resolved, multi-omics datasets from animal models with human systems represents a robust approach to identifying conserved pathways and prioritizing translational targets for AKI-induced lung injury.

In this study, we integrated transcriptomic and proteomic profiling across murine and porcine models of ischemia-reperfusion (IR) AKI, together with human lung organoids, to identify conserved early molecular events that mediate kidney-to-lung communication. To bridge the findings to human biology, we employed NHBE-derived lung organoids exposed to mtDAMP-stimulated macrophage-conditioned media. Using integrative multi-omics approaches (MOFA+, PROGENy, and DIABLO), we mapped the temporal molecular evolution of AKI-lung crosstalk and identified early vascular and structural targets with potential clinical relevance. To confirm our findings in humans, we also analyzed lung tissue from primary human organ donors. Together, our cross-species and multi-omics evidence demonstrates that AKI-induced lung injury involves an active, highly coordinated reprogramming of pulmonary gene regulation and metabolism.

## MATERIAL AND METHODS

### Analysis of Clinical Outcomes in AKI and Subsequent Lung Injury

This retrospective analysis used aggregate deidentified data from Epic Cosmos, a cross-institutional electronic health record (EHR) database, to describe clinical outcomes among patients with acute kidney injury (AKI) from January 1, 2016, to December 31, 2025. AKI was identified using the acute kidney failure code and all subcodes (ICD-10-CM: N17.*), and subsequent lung injury within 6 months of the initial AKI diagnosis was identified using SNOMED criteria. Queries identified the overall AKI cohort and the subgroup with subsequent lung injury (AKI-lung); hospital length of stay (LOS) was summarized from the reported AKI-alone and AKI-lung queries. Outcomes included all-cause mortality, mortality recorded with AKI as the preliminary cause of death, and hospital LOS. Mortality was summarized as aggregate counts and proportions with 95% confidence intervals. LOS was summarized using means, medians, and interquartile ranges; the reported comparison of mean LOS used a Welch two-sample t-test. Patient-level records were unavailable for this analysis. Consequently, patient-level multivariable regression, propensity score methods, and other analyses requiring individual covariate data could not be performed. Clinical comparisons were unadjusted and were interpreted as descriptive associations rather than independent or causal effects. Statistical tests were two-sided, with a significance threshold of p < 0.05. Analyses were conducted in R using RStudio (version 2025.05.1), with visualization using ggplot2.

### Animals and Bilateral IR-AKI Surgical Procedure

The IR-AKI mouse model was generated as described previously^3^. Briefly, mice were anesthetized with 2%–3% isoflurane, followed by a single subcutaneous injection of sustained-release buprenorphine (1 mg/kg; Wedgewood Connect, San Jose, CA) for analgesia. Through a midline abdominal incision, the kidneys were exposed, and both renal pedicles were carefully isolated. Ischemia was induced by applying vascular clamps to the pedicles for 30 min, after which the clamps were released to allow reperfusion. Core body temperature was maintained between 36.8°C and 37.2°C throughout the procedure. At 24 h post-reperfusion, animals were humanely euthanized for necropsy and downstream tissue harvesting.

The IR-AKI porcine model used female American Yorkshire pigs (4–5 months old, 60–70 kg), acclimated for at least 7–10 days prior to experiments. Animals were housed with free access to water and feed, with food withheld for ∼12 hours before surgery. Anesthesia was induced with ketamine (3 mg/kg), xylazine (3 mg/kg), and midazolam (0.4 mg/kg), followed by endotracheal intubation and maintenance on 2–3% isoflurane in oxygen (2 L/min). Pigs were mechanically ventilated with parameters adjusted to maintain physiological blood gases. Bilateral renal ischemia was induced by clamping both pedicles for 90 minutes, then reperfused for 4 hours. Sham controls underwent surgical exposure without clamping. Animals were euthanized for tissue collection. All procedures followed NIH guidelines and were approved by the University of South Florida Institutional Animal Care and Use Committees (IACUC protocol No.: 12005R for mice, 12931R for pigs).

### High-Performance Liquid Chromatography (HPLC) Measurement of Plasma Creatinine

Plasma creatinine was measured by HPLC following established protocols with minor modifications^3,19^. Briefly, 5 μl of plasma was protein-precipitated with 200 μl acetonitrile, vortexed, and centrifuged at 13,000 × g for 15 minutes at 4°C. The clear supernatant was dried under reduced pressure and reconstituted in 50 μl of mobile phase. Samples were analyzed on an Agilent 1260 Infinity II HPLC system using a Raptor C18 column (2.7 μm, 100 × 3.0 mm) with 5 mM sodium acetate as the mobile phase at a flow rate of 0.3 ml/min. Detection was at 225 nm, and the creatinine retention time was 1.67 ± 0.02 min.

### Generation of Lung Organoid

The primary Normal Human Bronchial Epithelial Cells (NHBEs) were a gift from Dr. Bailey (Nebraska Medical Center, Omaha, NE). The cells (P3) were cultured in a T-25 flask with PneumaCult^TM^-Ex Plus Medium (STEMCELL, Cambridge, MA) at 37^°^C in an atmosphere of 5% CO_2_. The medium was changed every two days. For three-dimensional (3D) organoid cultures, cells at 60% confluency were mixed with seeding medium and Corning Matrigel®, then plated as domes in a prewarmed 24-well plate according to the PneumaCult™ Airway Organoid Kit protocol. After polymerization, wells were overlaid with PneumaCult™ Airway Organoid Seeding Medium. Following 7 days of incubation, the culture medium was switched to PneumaCult™ Airway Organoid Differentiation Medium and changed every two days for an additional 28 days. Organoids were imaged using the EVOS XL imaging system (Life Technologies, USA).

### Preparation of mtDAMPs

Mitochondrial damage-associated molecular patterns (mtDAMPs) were prepared from human HK-2 cells. Mitochondria were isolated using a commercial isolation kit (Miltenyi Biotec) following the manufacturer’s instructions. The mitochondrial fraction was resuspended in DMEM and subjected to sonication on ice (30-second pulses alternating with 30-second rests, for 10 cycles) to release mitochondrial contents. Lysates were centrifuged sequentially at 12,000 × g for 10 minutes and then at 100,000 × g for 30 minutes at 4°C to remove membranes and debris. The supernatant containing soluble mtDAMPs was collected, and protein concentration was measured using the Pierce™ BCA Protein Assay Kit (Thermo Fisher Scientific).

### Immunocytochemical staining of Human lung organoid

Organoid harvesting, cleaning and fixation were performed as described elsewhere with modification ^20^. Briefly, the Matrigel-cell mixture was dissolved with cold Corning Cell Recovery Solution (Sigma), and organoids were collected by gentle centrifugation (70 × g, 4°C). After rinsing in ice-cold PBS, organoids were fixed in 4% paraformaldehyde for 45 minutes at 4°C, then washed with 0.1% PBS-Tween and blocked in organoid washing buffer (1 ml Triton X-100 and 2 g BSA per 1 L PBS) for 15 minutes. Organoids were incubated overnight at 4°C with primary antibody (1:100), washed, and then incubated overnight with Alexa Fluor 594 secondary antibody (1:500, Life Technologies). After final washes, 50 μl FocusClear (CelExplorer) was added for clearing and imaging. Imaging was performed on an Olympus FV1200 laser scanning confocal microscope (Olympus, Tokyo, Japan).

### RNA and protein sample preparation, sequencing and data processing

Total RNA was isolated from lung tissue using the RNeasy Mini Kit (QIAGEN, Hilden, Germany) according to the manufacturer’s instructions. Briefly, approximately 30 mg of tissue was homogenized in RLT buffer with β-mercaptoethanol, and RNA was purified via spin columns and assessed for concentration and purity by NanoDrop™ spectrophotometry. Purified RNA was stored at −80°C and submitted for sequencing. For proteomics, lung tissue was flash-frozen in liquid nitrogen and stored at −80°C before analysis.

The RNA samples were delivered to Novogene (Beaverton, OR) for sample quality control, library preparation, and high-throughput sequencing. Raw FASTQ files underwent quality control using FastQC (v0.12.1), and adapter sequences, poly-N stretches, and low-quality bases were trimmed using fastp. Cleaned, high quality paired-end reads were aligned to the reference genome (Homo_sapiens.GRCh38.114 for human; Sus_scrofa.Sscrofa11.1.114 for pig; Mus_musculus.GRCm39.114 for mice) using STAR (v2.7.10b) with default parameters. Read counts per gene were quantified using featureCounts (v2.0.6) against Ensembl gene annotations. Gene-level count matrices were imported into R (4.5.1) for differential expression analysis using DESeq2 (1.50.2).

Flash-frozen samples were shipped on dry ice to Biognosys (Newton, MA) for high-throughput quantitative proteomics. Tissue samples were lysed, denatured, reduced, and alkylated by Biognosys with their protocol. The DIA mass spectrometric data were acquired using Evosep One liquid chromatography system connected to a Bruker Daltonics timeTOF HT mass spectrometer equipped with a Captive Spray II ion source at their facility and analyzed using Spectronaut software (Biognosys, version 20.5) with UniProt .fasta database (Hono sapiens; Sus scrofa; Mus musculus, 2026-01-01). Log2-transformed protein intensities were analyzed in R using the limma package (3.66).

### MOFA+ Analysis

Six omics datasets were prepared for multi-omics factor analysis (MOFA+). Proteomics data were log2-transformed for normalization. RNA-seq data underwent variance-stabilizing transformation (VST) using DESeq2. The top 3,000 variable features (proteomics) and top 5,000 (RNA-seq) per dataset were selected based on variance, excluding features with zero or missing variance.

Multi-omics factor analysis was performed using the MOFA2 R package (version 1.20.2) with the mofapy2 Python backend (version installed via reticulate). Since the three experimental models comprised unequal sample sizes (pig: n=10, mouse: n=10, human organoid: n=8), a single-group model was constructed by padding missing samples with NA values across views. MOFA+ natively handles missing data through a variational Bayesian expectation-maximization algorithm, allowing features measured in only a subset of samples to contribute to factor inference without imputation. The MOFA+ model was configured with the following parameters: number of factors = 10, scale views = TRUE (to account for different variance scales across omics layers and species), convergence_mode = “medium”, maximum iterations = 1,000, and random seed = 42 for reproducibility. Sample metadata including species, experimental model, condition (AKI vs Control), and timepoint (4hr vs 24hr) were annotated to the MOFA+ object using the samples_metadata function.

The proportion of variance explained by each factor in each omics view (R²) was calculated and visualized as a heatmap. Factors were interpreted based on the variance they explained per view, the mean factor values per condition and model, and the magnitude of AKI versus control separation (delta = mean AKI – mean Control). Factors were classified as model-specific if they explained >10% variance in only one dataset, or as cross-species conserved if they explained >2% variance across two or more datasets. By these criteria, Factor 1 was mouse-specific, Factor 2 was human organoid-specific, Factor 3 was pig-specific, and Factors 5, 6, and 7 were cross-species conserved.

For cross-species conserved Factors 5, 6, and 7, the top 100 genes (by absolute weight) were extracted from each view. Gene symbols were standardized and analyzed for up- and down-regulation in AKI. Over-representation analysis (ORA) was conducted using clusterProfiler and the human gene annotation database (org.Hs.eg.db), with all annotated features (n=11,169) as the background. Significant Gene Ontology (GO) biological process terms were defined as having BH-adjusted p < 0.05 and q < 0.2, and results were visualized in bubble plots showing gene counts and adjusted p-values.

### Pathway RespOnsive GENes (PROGENy) pathway activity and Transcription factor activity analysis

Pathway activity was estimated using the PROGENy R package (v1.32.0), computing activity scores for 14 pathways across RNA and protein datasets. Statistical significance between AKI and control was assessed using the Wilcoxon rank-sum test. Transcription factor activity was inferred using decoupleR (v2.16.0) with the multivariate linear model approach and the CollecTRI regulon database (43,159 TF-target interactions, 1,186 TFs). The minimum regulon size was set to 5 targets. For each model, TF activity delta scores were calculated as the difference in mean activity between AKI and control. Conserved TFs were those ranking in the top 30 by absolute delta score in at least two models.

### Data Integration Analysis for Biomarker discovery using Latent components in Omics studies (DIABLO) analysis

Multi-omics biomarker discovery was conducted using DIABLO (mixOmics R package, v6.34.0). Separate DIABLO models were constructed for each experimental system, integrating paired RNA and protein data. Input features included significant differentially expressed proteins (DEPs, |log2FC| > 1, raw p < 0.05) and the most variable RNA features (top 200 for pig and mouse; top 50 for human organoid, due to sample size). The design matrix off-diagonal value was set at 0.1 to balance outcome discrimination and block correlation. Feature selection (keepX) was optimized by leave-one-out cross-validation for pig and mouse models and fixed at 5 per block for the human organoid model (n=8). Classification accuracy was evaluated by leave-one-out cross-validation error rate.

### Taqman Real-time PCR (RT-PCR)

Total RNA was isolated from tissue homogenates from primary human donor lungs using TRIzol^®^ reagent. One-step quantitative reverse transcription-PCR was performed using the Taqman RNA-to-Ct 1-Step Kit (Applied Biosystems, Foster City, CA) on an automated real-time PCR detection system. Target gene transcript levels were quantified using pre-designed TaqMan Gene Expression Assays (Applied Biosystems, GAPDH: Hs02786624_g1; FOXF1: Hs00230962_m1; ATF6: Hs00232586_m1; CLOCK: Hs04976593_s1). Relative expression changes were calculated using the 2^−ΔΔCt^ method.

### Data Analysis and Statistics

Statistical analyses were performed using GraphPad Prism version 11.0.1 (GraphPad Software, San Diego, CA). P-values for volcano plots were adjusted by the Benjamini-Hochberg method to control the false discovery rate (FDR). Data are reported as mean ± standard error (SE).

## RESULTS

### Aggregate clinical outcomes in the AKI cohort and AKI-lung subgroup

At the time of the study query in May 2026, Epic Cosmos included 301,125,838 patients Between January 1, 2016, and December 31, 2025, 13,815,092 patients were identified with AKI. In this overall AKI cohort, 3,515,757/13,815,092 patients (25.45%, 95% CI: 25.43%–25.47%) had recorded all-cause mortality. In the subgroup with lung injury within 6 months after AKI, 7,284/19,862 patients (36.67%, 95% CI: 36.00%–37.34%) had recorded all-cause mortality. Mortality recorded with AKI as (25.45% vs. 36.67%, χ² = 1315.9, p < 0.001) (Figure 1A). For the preliminary cause of death occurred in 7,079/13,815,092 patients (0.05%, 95% CI: 0.049%–0.051%) in the overall AKI cohort and 17/19,862 patients (0.09%, 95% CI: 0.05%–0.14%) in the AKI-lung group, also demonstrating a statistically significant difference (χ² = 4.56, p = 0.033) subgroup (Figure 1B). These aggregate estimates describe the observed outcomes without adjustment for patient characteristics.

**Figure 1.**
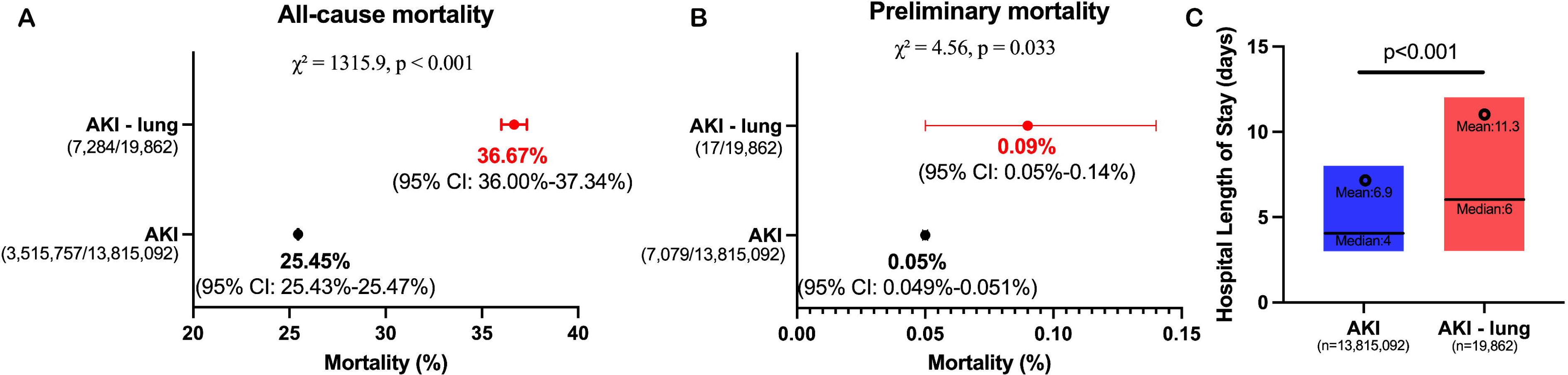
Aggregate clinical outcomes among patients with AKI and subsequent lung injury. (A) Recorded all-cause mortality in the overall AKI cohort and the subgroup with lung injury within 6 months after AKI. (B) Mortality recorded with AKI as the preliminary cause of death in the same queries. (C) Hospital length of stay (LOS) in the reported AKI-alone and AKI-lung queries. Mortality was summarized using aggregate counts and proportions. LOS was summarized using means, medians, and interquartile ranges; the reported comparison of means used a two-sided Welch two-sample t-test with a significance threshold of p < 0.05. All clinical comparisons were unadjusted. Analyses were conducted in R using RStudio (version 2025.05.1); this figure was visualized using GraphPad Prism (version 11.0.1).

Hospital length of stay (LOS) was also longer among patients in the AKI-lung group. Median LOS was higher in the AKI-lung group (6 days, IQR: 3–12) compared with the AKI alone group (4 days, IQR: 3–8), with a wider interquartile range indicating greater variability. Mean LOS was also higher (11.3 vs. 6.9 days) and differed significantly between groups (Welch two-sample t-test, p < 0.001) (Figure 1C). The greater divergence between mean and median in the AKI-lung group suggests a more right-skewed distribution, consistent with prolonged hospitalizations in a subset of patients. These unadjusted findings describe longer LOS in the AKI-lung query; they do not establish an independent effect of lung injury on hospitalization duration.

### IR-AKI induces remote organ lung injury in mice

To investigate the molecular connection between acute kidney injury and distant lung dysfunction, we first characterized the impact of IR-AKI in a murine model. AKI was confirmed by a significant elevation in plasma creatinine at 24 hours post-reperfusion (Figure 2A). Principal component analysis (PCA) showed distinct clustering by experimental group, indicating that IR-AKI induces robust shifts in both gene (Figure 2B) and protein (Figure 2C) expression within the lung. Integration of transcriptomic and proteomic profiles revealed a multi-level response to IR-AKI. RNA-seq identified 2,384 differentially expressed genes (DEGs; 605 upregulated, 1,779 downregulated; Supplementary file S1, Figure 2D), while proteomic analysis revealed 80 DEPs (45 upregulated, 35 downregulated; Supplementary file S1, Figure 2E). Spearman correlation of 9,310 matched features demonstrated a significant positive correlation (r = 0.316), consistent with known proteogenomic trends in acute injury models (Figure 2F). While many features were discordant, likely reflecting post-transcriptional regulation, a subset of 45 genes showed robust directional concordance (*P_adj_* < 0.05, |LFC| > 0.5 at both levels). Integrated heatmap (Figure 2G) analysis confirmed that the pulmonary response to remote kidney injury was driven by transcriptional activation and translation of inflammatory effectors. Gene Ontology (GO) enrichment showed that upregulated genes highlighted pathways involved in structural remodeling and metabolic signaling, with PI3K/Akt signal transduction (*P_adj_* = 6.12 × 10⁻⁶) most significantly enriched. Additional terms included collagen metabolism, smooth muscle cell proliferation, and cellular responses to insulin, indicating a multifaceted response involving extracellular matrix reorganization and altered metabolism (Figure 2H & 2I).

**Figure 2.**
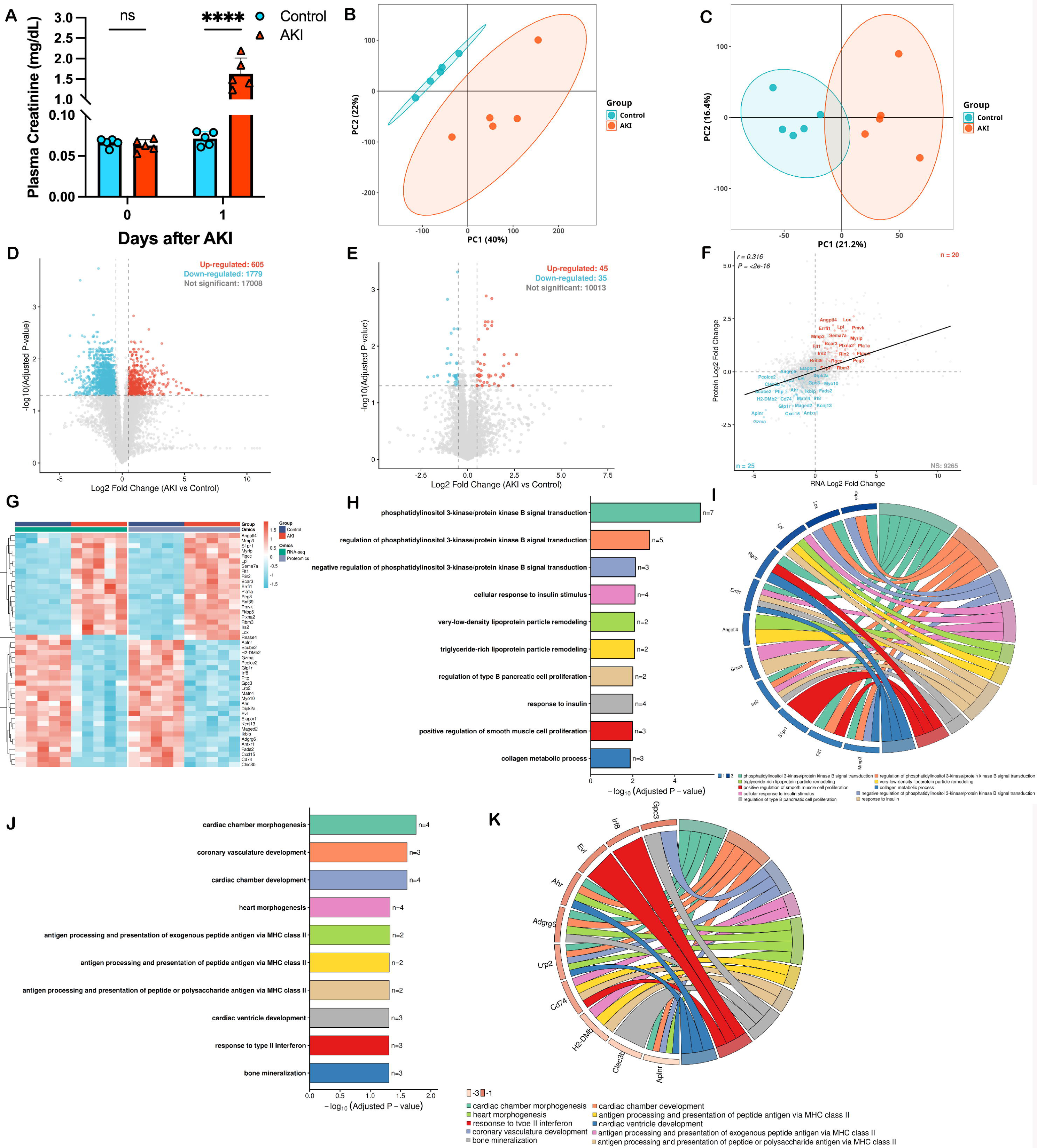
Proteogenomic Characterization of Mouse Lung Response to IR-AKI. (A) Plasma creatinine levels dramatically increased in mice following IR-AKI surgery (n=5 per group; **** p < 0.0001, unpaired t-test). (B-C) PCA of RNA-seq (B) and proteomics (C) revealed clear separation between control and AKI lung samples. (D-E). Volcano plots of differentially expressed genes (DEGs; D) and proteins (DEPs; E) in lung tissue after AKI (|LFC| > 1, *P_adj_* <0.05). (F) Scatter plot of log2 fold changes for matched transcripts and proteins; a core signature of 20 features was upregulated, and 25 were downregulated at both the transcript and protein level. (G) Integrated heatmap of significant concordant features across RNA and protein datasets. (H) Top 10 enriched Gene Ontology (GO) biological processes among upregulated genes. (I) Chord plot showing connections between the top 20 upregulated genes and their associated GO terms. (J) Top 10 enriched GO terms among downregulated genes, highlighting suppression of vascular and immune pathways. (K) Chord plot illustrating interactions between core downregulated hub genes and structural maintenance pathways.

Enrichment analysis of downregulated genes revealed suppression of programs required for vascular and structural maintenance, such as cardiac chamber morphogenesis (*P_adj_* = 0.0175) and coronary vasculature development (*P_adj_* = 0.0247), suggesting disruption of genetic programs essential for pulmonary vascular integrity (Figure 2J). Further analysis identified six genes bridging structural maintenance and immune signaling: Aplnr, Lrp2 (vascular stability), Adgrg6, Ahr (structural integrity), and Cd74, H2-DMb2 (immune surveillance). The coordinated loss of vascular and immune genes indicates that AKI-induced lung injury severity is influenced by both activation of inflammatory pathways and simultaneous failure of homeostatic networks (Figure 2K).

### Acute renal injury triggers early proteomic shift prior to transcriptional divergence in pig model

To examine the immediate pulmonary response to AKI in a large-animal model, we induced bilateral renal ischemia for 90 minutes followed by 4 hours of reperfusion in pigs. AKI was confirmed by a significant elevation in plasma creatinine at 4 hours post-reperfusion (Figure 3A). PCA of lung transcriptomics showed no clear separation between AKI and sham groups at this early time point (Figure 3B), indicating that the transcriptome was largely unchanged. In contrast, proteomic profiling revealed a distinct molecular shift in the AKI group, as shown by clear group separation in PCA (Figure 3C).

**Figure 3.**
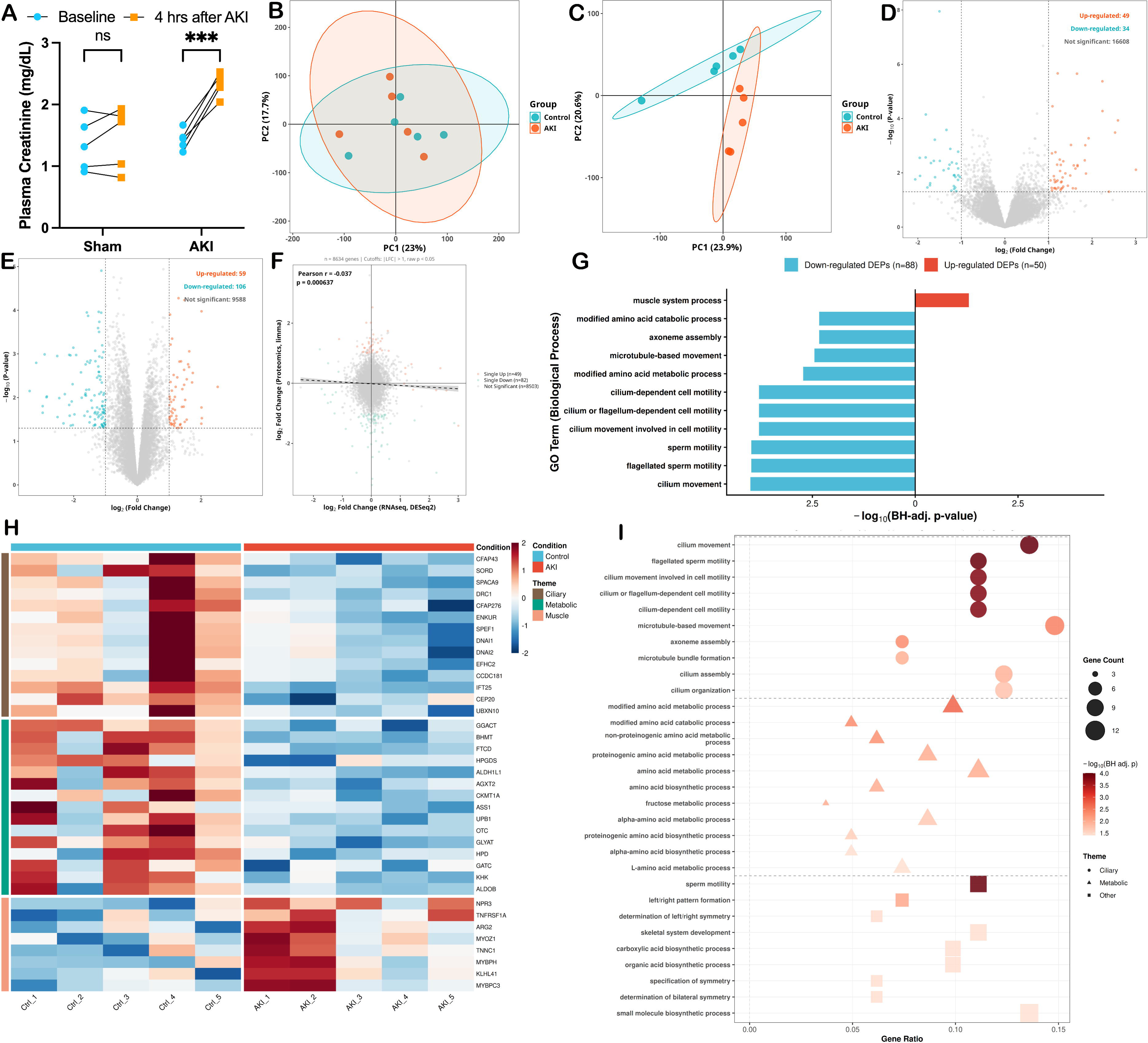
Proteogenomic Characterization of Pig Lung with 90 min IR-AKI followed by 4 hrs reperfusion. (A) Plasma creatinine levels significantly increased post-AKI (n = 5 per group; ***p < 0.001, paired t-test). RNA-seq PCA showed no clear separation between control and AKI lung samples. (C) Proteomic PCA revealed distinct separation between groups. (D, E) Volcano plots of differentially expressed genes (DEGs; D) and proteins (DEPs; E) in lung tissue (|LFC| > 1, p < 0.05). (F) Correlation analysis of log2 fold changes for matched transcripts and proteins (Pearson r = –0.06). (G) Top 10 upregulated and downregulated GO biological processes among concordant features. (H) Heatmap of key DEPs involved in these GO terms. (I) Network analysis showing translational recovery and metabolic suppression clusters.

Transcriptomic analysis identified only 83 DEGs; 49 upregulated, 34 downregulated; Figure 3D), suggesting that early lung responses to AKI are limited at the transcriptional level. Proteomics, however, revealed 164 DEPs; 58 upregulated, 106 downregulated; Figure 3E). Although none remained significant after BH correction (likely due to the early time point and model variability), they represent high-confidence candidates for further study (Supplement S2). Correlation analysis of 8,634 matched features showed a weak but significant negative correlation between transcriptomic and proteomic changes (r = – 0.037, p = 0.000637; Figure 3F), indicating that early lung responses are predominantly regulated post-transcriptionally.

GO enrichment of concordant upregulated proteins highlighted pathways related to ribosome biogenesis and rRNA processing, while concordant downregulated proteins were enriched for mitochondrial energy and TCA cycle processes (Figure 3G). This suggests a metabolic shift away from oxidative phosphorylation toward enhanced translational capacity. Targeted analysis of genome-wide proteomic and translational hubs revealed distinct regulatory patterns: translational regulators (e.g., BMS1) increased more at the protein than transcript level. At the same time, metabolic enzymes (e.g., PDHA1) showed protein depletion despite unchanged mRNA (Figure 3H). Network analysis identified two main modules: a translational recovery cluster (BMS1, NPM1, UTP4) linked to ribosome biogenesis, and a metabolic suppression cluster (PDHA1, ACADM) associated with downregulated energy metabolism (Figure 3I).

### Cross species proteomic comparison of different reperfusion stages

AKI-induced lung injury evolves over time. To explore the temporal evolution of AKI-induced lung injury, we compared systemic molecular responses in pig lung tissue at 4 hours post-reperfusion and mouse lung tissue at 24 hours post-reperfusion. Due to limited changes in pig RNA-seq data, proteomic data with a relaxed fold-change threshold (|Log2FC| > 0.5, p < 0.05) were used for cross-species comparisons. In total, 267 upregulated and 375 downregulated proteins were identified in pig lung tissue, while the mouse model exhibited 341 upregulated and 237 downregulated proteins (Figure 4A & 4B). Overlap between species was limited, with only 17 conserved upregulated and 5 conserved downregulated proteins, consistent with distinct biological phases in each model. Cross-species correlation of protein Log_2_FC values for 8,145 shared proteins indicating that the pig and mouse protein responses were essentially uncorrelated (Figure 4C). Examining the top DEPs highlighted the temporal shift in biological response: in pigs at 4 hours, downregulated proteins were primarily structural and metabolic, whereas in mice at 24 hours, upregulated proteins included immune effectors and inflammatory mediators (Figure 4D). GO enrichment of pig downregulated proteins revealed early ciliary dysfunction and disrupted amino acid metabolism. Meanwhile, mouse upregulated proteins were enriched for acute inflammatory response, leukocyte migration, and bacterial defense (hallmarks of established injury) (Figure 4E). Importantly, RNA-protein concordance increased from pig (r = –0.06) to mouse (r = 0.447), illustrating a shift from post-translational to transcriptional regulation as the injury progresses over time (Figure 4F).

**Figure 4.**
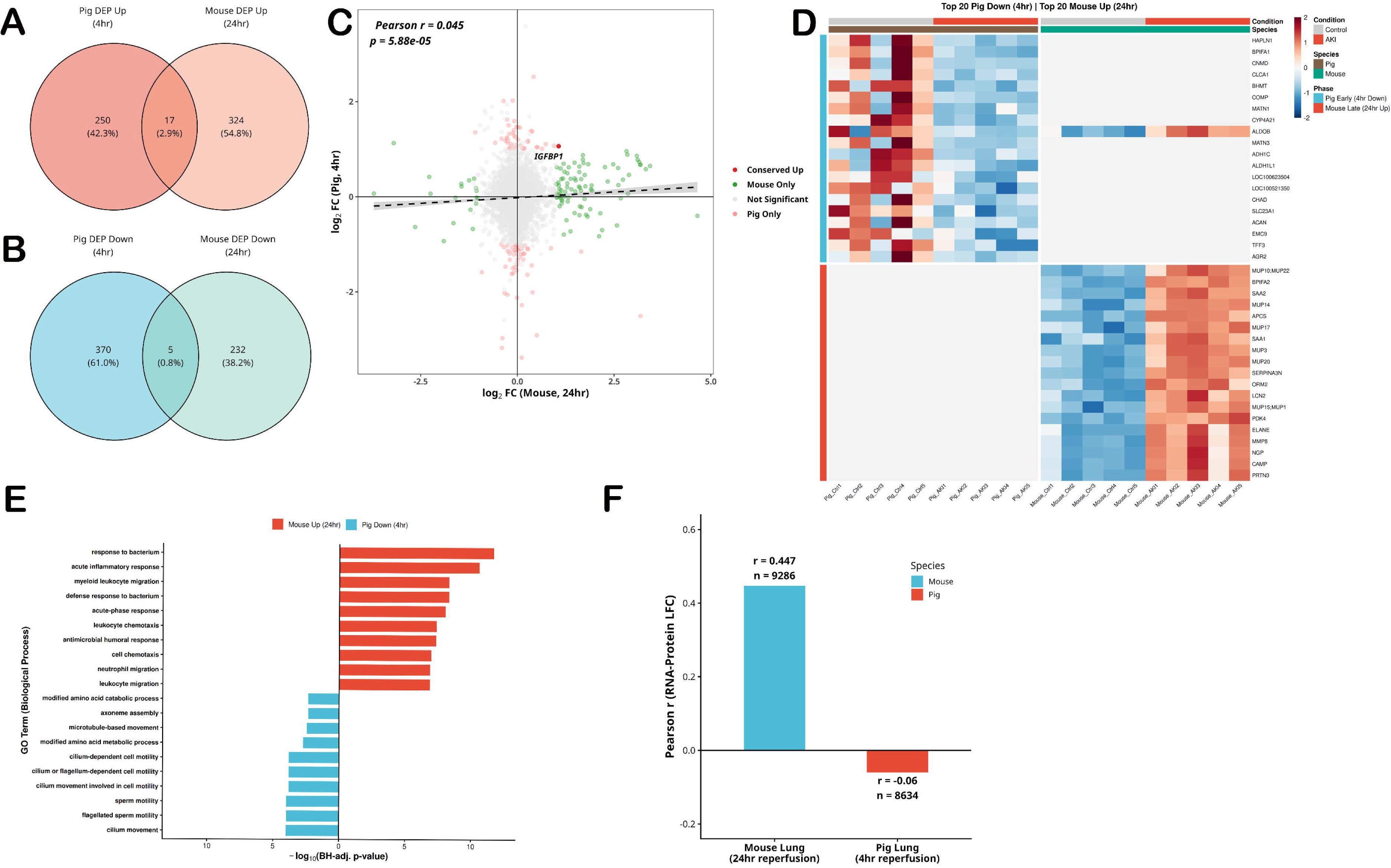
Cross-species proteomic integration reveals temporal phases of IR-AKI-induced lung injury. (A) Venn diagram of upregulated DEPs in pig and mouse lung tissue. (B) Venn diagram of downregulated DEPs in both species. (C) Scatter plot of protein Log2FC across all shared proteins (n = 8,145); linear regression with 95% confidence interval. (D) Z-score heatmap of top 20 pig downregulated (blue) and mouse upregulated (red) proteins. (E) Bar plot of top 10 GO biological processes enriched among pig and mouse DEPs (–log10 BH-adjusted p-value). (F) Bar plot comparing RNA-protein Log2FC concordance (r) between pig and mouse lung, illustrating the temporal regulatory shift.

### mtDAMPs treatment revealed a recovery response in NHBE lung organoid model

Building on previous findings that AKI promotes lung injury via release of mitochondrial damage-associated molecular patterns (mtDAMPs), we tested the effects of mtDAMPs on human lung organoids. mtDAMPs were isolated from HK-2 cells and used to treat human lung organoids via conditioned media from mtDAMP-stimulated macrophages (Figure 5A). Organoid differentiation and structure were confirmed by brightfield imaging and immunocytochemical staining for ZO-1 (cell boundaries), MUC5AC (goblet cells), and acetylated tubulin (ciliated cells) (Figure 5B, 5C).

**Figure 5.**
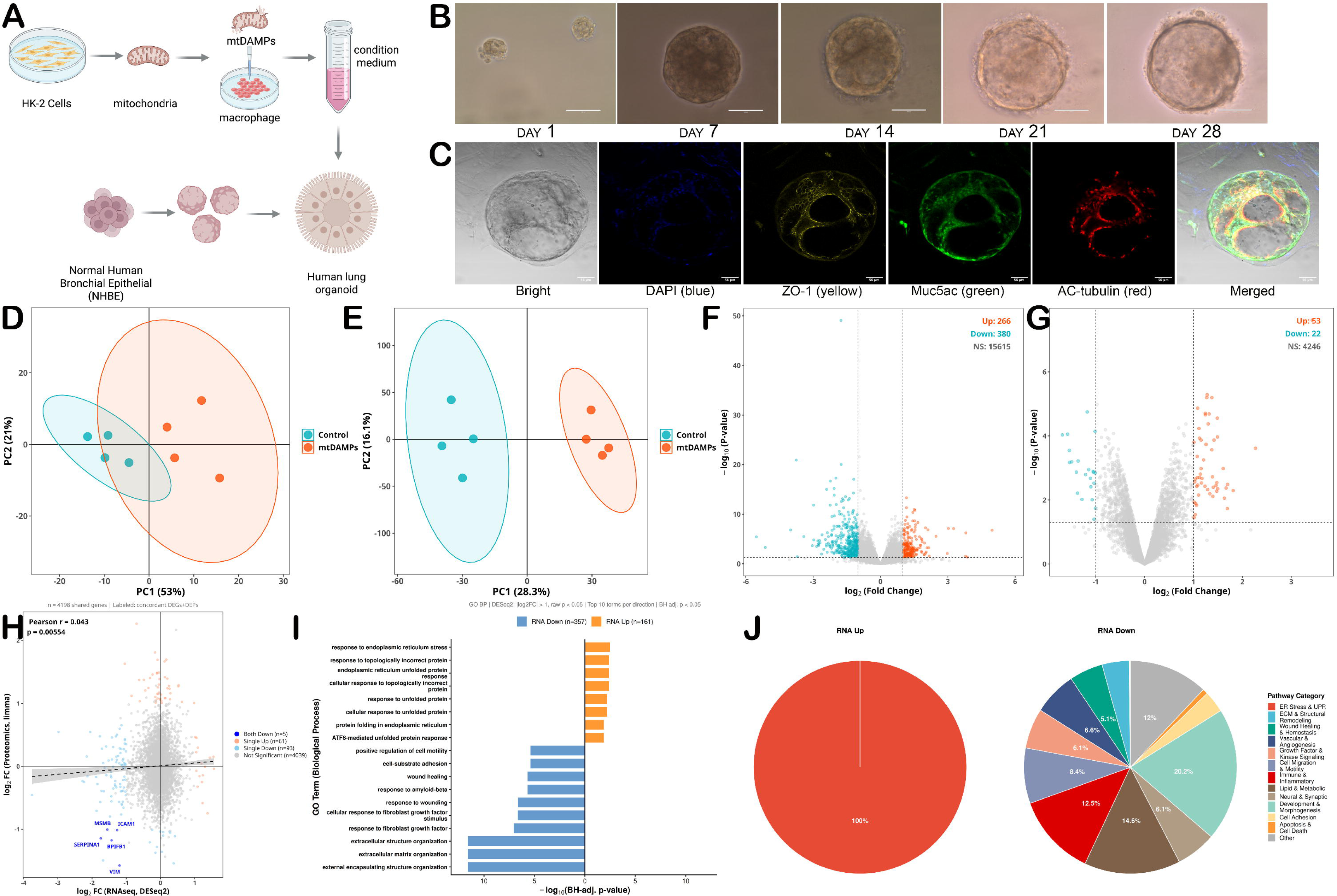
Human lung organoid multiomics response to mtDAMPs-conditioned macrophage media. (**A**) Schematic of experimental workflow: mtDAMPs from HK-2 cells were used to stimulate macrophages, and conditioned media was applied to NHBE-derived lung organoids for 24 hours. (B) Brightfield images showing organoid differentiation over 28 days. (C) Immunofluorescence staining for ZO-1 (yellow), MUC5AC (green), and acetylated tubulin (red). (D) PCA of RNA-seq data; (E) PCA of log2-transformed proteomics data. (F) Volcano plot of DEGs (|log2FC| > 1, p < 0.05). (G) Volcano plot of DEPs (|log2FC| > 1, p < 0.05). (H) Scatter plot of RNA-protein log2FC for shared genes. (I) Bar plot of top 10 GO biological processes for up- and downregulated genes. (J) Pie chart of GO pathway categories in mtDAMP-treated organoids.

After 24 hours of exposure, RNA-seq and proteomic analyses were performed. PCA demonstrated clear separation between mtDAMP-treated and control organoids at both the transcriptomic (Figure 5D) and proteomic (Figure 5E) levels. A total of 266 upregulated and 380 downregulated genes were identified by RNA-seq (|log_2_FC| > 1, p < 0.05), while proteomics identified 74 upregulated and 30 downregulated proteins (Figure 5F, 5G). RNA-protein concordance was low (r = 0.035, n = 4,198 shared genes), with only 5 genes concordantly downregulated and none concordantly upregulated at both levels (Figure 5H). This decoupling, along with the predominance of downregulated transcripts, suggests that the organoids had entered a resolution or tissue repair phase by 24 hours post-treatment.

GO analysis of upregulated genes highlighted enrichment for endoplasmic reticulum (ER) stress and unfolded protein response (UPR) pathways (Supplementary S3), including ATF6-mediated UPR and ER protein folding. In contrast, downregulated genes were enriched for extracellular matrix organization, wound healing, fibroblast growth factor signaling, and cell motility, indicating coordinated suppression of tissue remodeling and active resolution (Figure 5I, 5J).

Together, these findings indicate that mtDAMP-conditioned macrophage media induces a transient but coordinated injury resolution response in human lung organoids, characterized by sustained ER stress and suppression of tissue remodeling pathways.

### Model-specific and conserved factors identified by Multi-Omics Factor Analysis v2 (MOFA+)

To identify shared and model-specific molecular signatures across species and experimental systems, we applied MOFA+ to integrate transcriptomic and proteomic data from pig, mouse, and human organoid models. The analysis revealed three model-specific factors: mouse-specific inflammatory response (Factor 1), human organoid-specific signal (Factor 2), and pig-specific early structural damage (Factor 3). Importantly, two factors—Factor 6 and Factor 7—emerged as cross-species conserved signals (Figure 6A).

**Figure 6.**
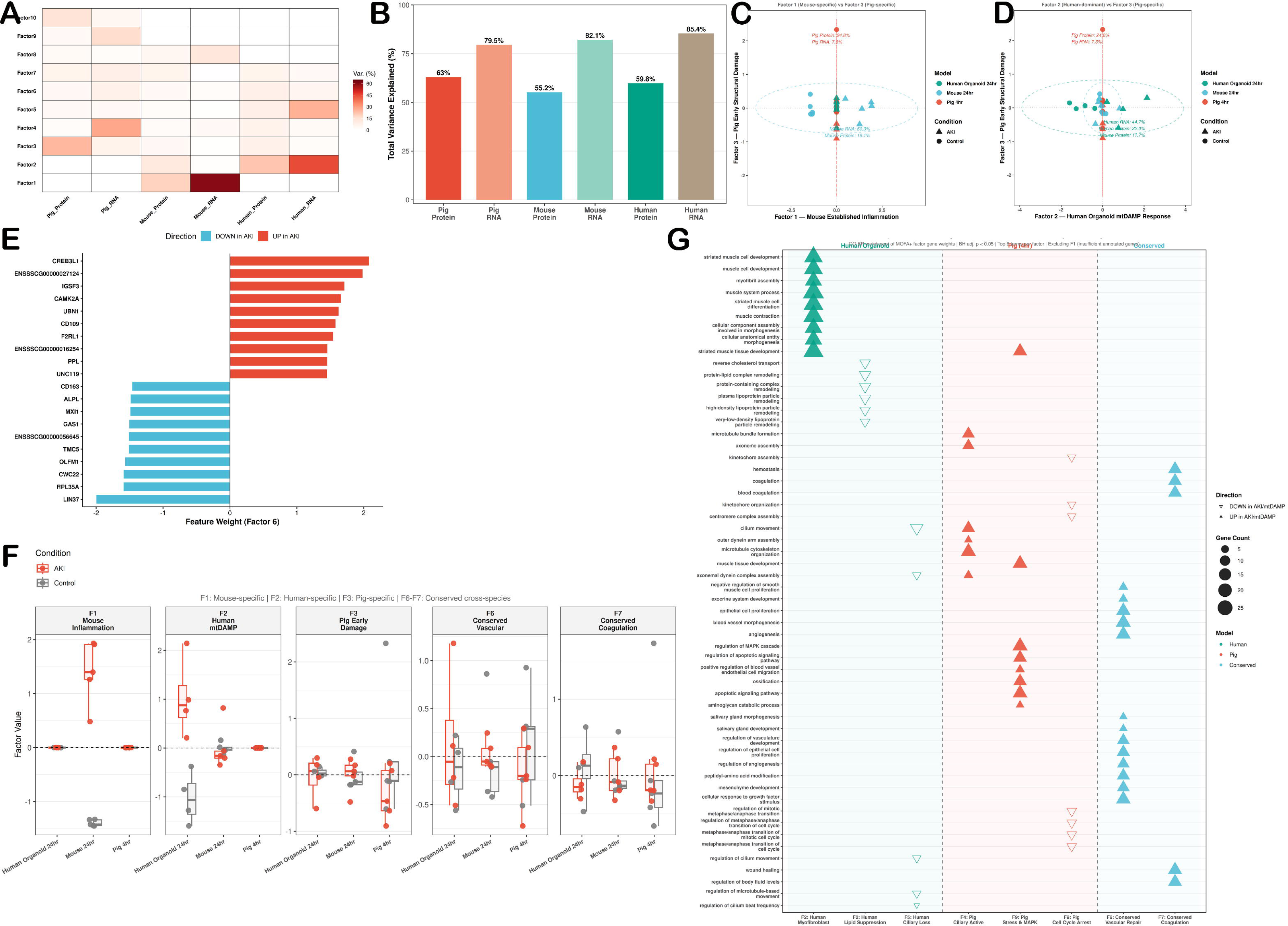
MOFA+ cross-species multi-omics integration, MOFA+ was trained on top variable features from each dataset (3000 proteins, 5000 RNA features per species) with 10 factors. (A) Heatmap of variance explained (R²) by each MOFA+ factor across all omics datasets. (B) Total variance explained per dataset. (C) Factor scatter plot—Factor 3 (pig) vs. Factor 1 (mouse); each point represents a sample. (D) Factor 2 (human organoid) vs. Factor 3 (pig), showing model-specific signals. (E) Top weighted genes driving Factor 6 in the pig protein dataset. (F) Factor values across all samples for key factors. (G) Bubble plot of significant GO biological process terms for top weighted genes in MOFA+ factors, highlighting model-specific and conserved biological programs.

Variance explained by each factor was presented in Figure 6B. Factor scores in the scatterplots showed clear separation of mouse samples (Factor 1), pig samples (Factor 3), and human organoids (Factor 2), while Factor 6 was the most conserved signal across all omics layers (Figure 6C–F). GO enrichment analysis of the top weighted genes in Factor 6 showed significant overrepresentation of vascular and epithelial remodeling pathways, indicating that vascular repair and epithelial regeneration are core responses to AKI-induced lung injury across species (Figure 6G). Factor 7 was enriched for blood coagulation, hemostasis, regulation of body fluid levels, and wound healing, highlighting hemostatic dysregulation as another universally conserved feature. Complete factor weights, variance explained values, and GO enrichment results are provided in Supplementary S4. Together, these results suggest that despite differences in species, timing, and injury models, vascular and epithelial repair and coagulation dysregulation represent conserved molecular responses to AKI-induced lung injury.

### Upstream regulatory mechanisms were identified using Pathway RespOnsive GENes (PROGENy) pathway activity analysis, decoupleR and CollecTRI

To dissect upstream regulatory pathways and transcription factors involved in AKI-induced lung injury across species, we inferred pathway activity using PROGENy and estimated transcription factor activity using decoupleR with the CollecTRI regulon database. PROGENy analysis of RNA data revealed distinct pathway activation patterns in each model. In the mouse 24-hour model, NF-κB, TNF-α, VEGF, hypoxia, JAK-STAT, p53, androgen, and TRAIL pathways showed significant activation, consistent with a robust inflammatory and stress response (Figure 7A). No pathways reached statistical significance in the pig model at 4 hours, reflecting the early, pre-transcriptional state. Proteomic PROGENy analysis supported these findings, with directional concordance for key pathways such as MAPK, NF-κB, and TNF-α (Figure 7B & 7C).

**Figure 7.**
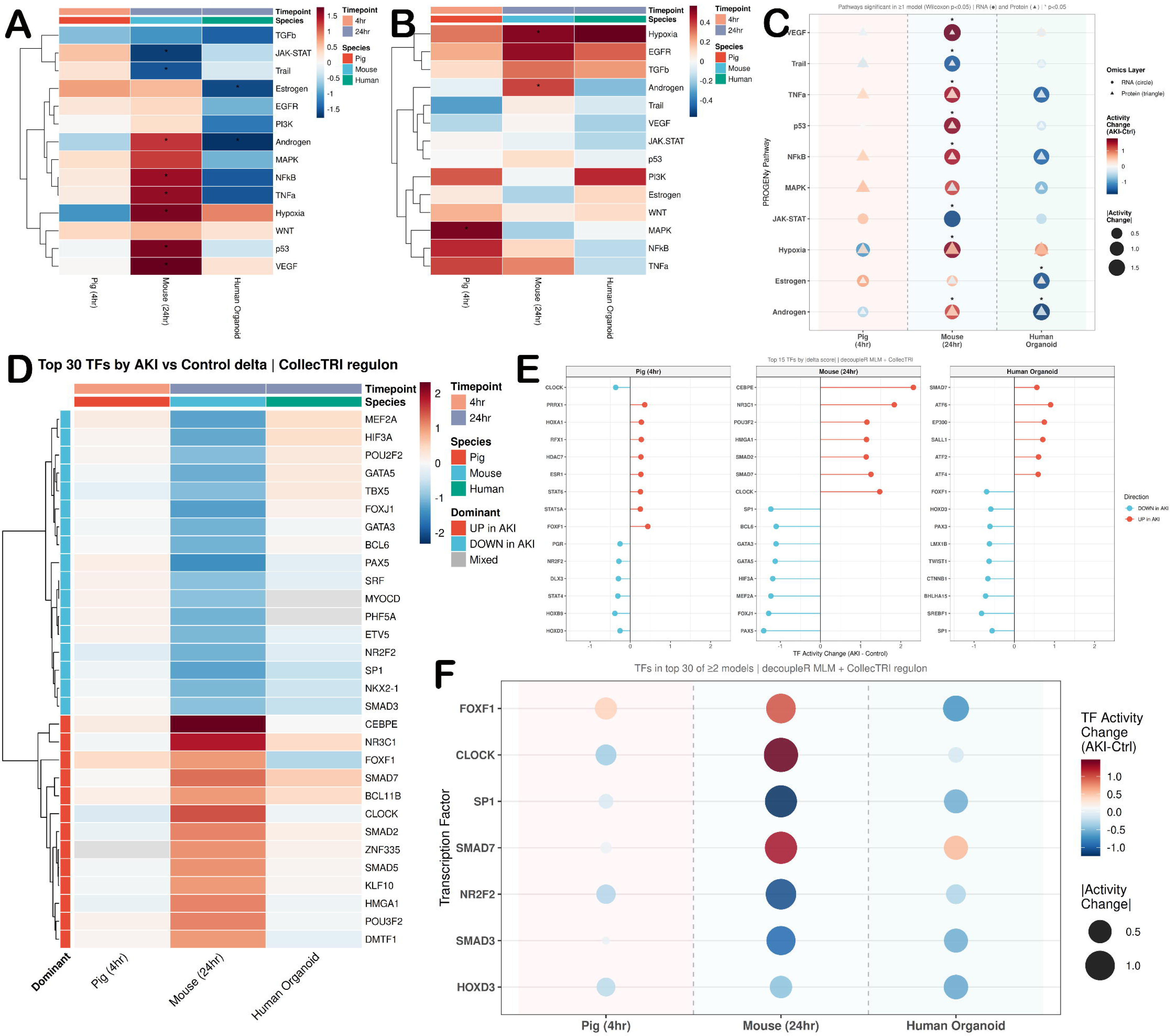
Cross-species pathway and transcription factor activity in AKI-induced lung injury. (A) Heatmap of PROGENy pathway activity z-scores (difference between AKI and control) across pig (4 hr), mouse (24 hr), and human organoid models using RNA-seq data; *p < 0.05, Wilcoxon test. (B) PROGENy pathway activity heatmap from proteomics data. (C) Bubble plot of significant PROGENy pathways across models and omics layers. (D) Heatmap of TF activity changes for the top 30 TFs per model, estimated with decoupleR MLM and the CollecTRI regulon. (E) Lollipop plots of the top 15 TFs by absolute activity delta per model. (F) Bubble plot of conserved TFs present in the top 30 in at least two models.

Transcription factor (TF) activity analysis using decoupleR and CollecTRI identified hundreds of TFs per model. In the pig model, top activated TFs included FOXF1, PRRX1, and HOXA1, associated with mesenchymal and epithelial identity maintenance (Figure 7D & 7E). The mouse model showed activation of CEBPE, NR3C1, CLOCK, SMAD7, and POU3F2, indicating inflammatory and circadian regulation. In human organoids, ATF6, EP300, ATF2, and ATF4 were most strongly activated, aligning with the ER stress signature seen in GO analysis. Among the top 30 TFs per model, seven were conserved across at least two models: FOXF1, CLOCK, NR2F2, HOXD3, SMAD7, SP1, and SMAD3 (Figure 7F). The conservation of SMAD3 and SMAD7—both TGF-β pathway regulators—suggests TGF-β signaling modulation as a fundamental feature of the AKI-lung axis across species.

### Multi-omics biomarker signatures were identified using Data Integration Analysis for Biomarker discovery using Laten components in Omics studies (DIABLO) analysis

To identify phase-specific biomarkers of AKI-induced lung injury in each experimental system, we applied DIABLO for integrated multi-omics analysis of paired RNA and protein data. Separate DIABLO models were constructed for human organoids, mouse, and pig datasets (Figure 8A). In the human organoid model, DIABLO selected FAP, POSTN, DKK3, COMP, and APOE as the top negatively weighted RNA biomarkers, reflecting suppression of extracellular matrix (ECM) and tissue remodeling pathways. At the protein level, CTSG and PNPLA2 were most strongly associated with AKI, indicative of residual inflammatory and lipid metabolic activity (Figure 8B & 8C), For the mouse model at 24 hours, the top RNA biomarkers included COLQ, SLC38A5, and APLN (negatively associated with AKI), while BPIFA2 and LOX were the leading protein biomarkers (positively associated), consistent with established-phase inflammatory and fibrotic remodeling (Figure 8B, 8D). In the pig model at 4 hours, Component 1 selected 15 RNA and 10 protein features. Notable discriminating proteins included IFT25 (linked to ciliary dysfunction), C19ORF12, NDST1, and PHC1, while at the RNA level, THBD, PPBP, and RGS5 were prominent (Figure 8B, 8E). The DIABLO model demonstrated the classification accuracy by leave-one-out cross-validation in each species (Figure 8F). Some pig-specific transcripts lacked annotation due to limitations in the Sus scrofa genome. Notably, no biomarkers were shared across models (Figure 8G), indicating that each experimental system captures a phase-specific molecular signature of AKI-induced lung injury. This finding aligns with the temporal evolution identified by MOFA+ and the regulatory differences highlighted by PROGENy and decoupleR, collectively indicating that AKI-lung injury progresses through distinct, dynamic molecular phases in each system.

**Figure 8.**
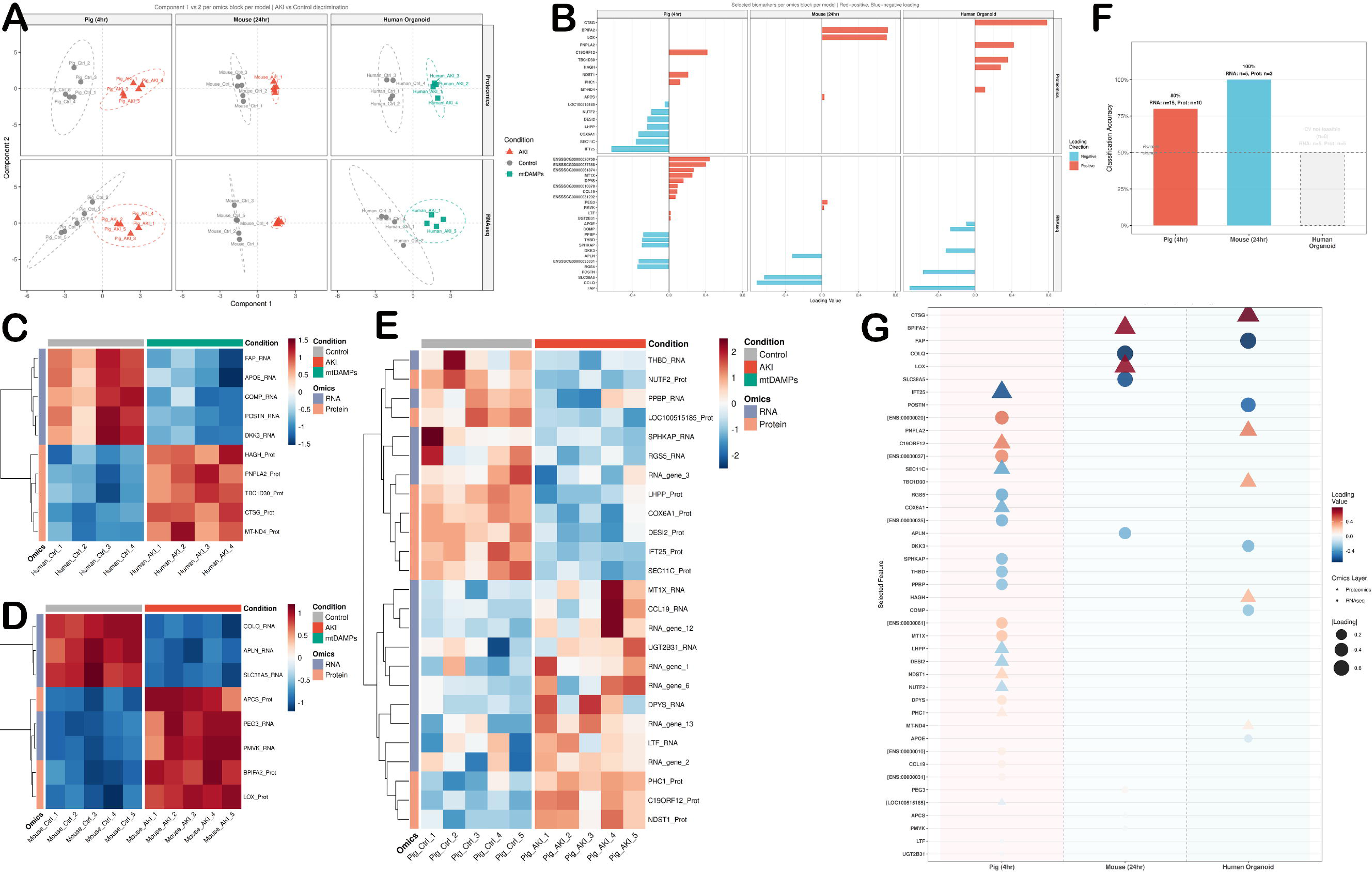
DIABLO multi-omics biomarker discovery across AKI-lung injury models. (A) DIABLO sample plots: Component 1 vs. Component 2 for RNA and protein blocks per model. (B) Feature loading plots for Component 1 per omics block and model. (C–E) Z-score heatmaps of DIABLO-selected biomarkers for human organoid (C), mouse (D), and pig (E) models; rows = features, columns = samples. (F) Leave-one-out cross-validation accuracy for pig (80%) and mouse (100%) models; the human organoid model was excluded from cross-validation due to small sample size. (G) Bubble plot of all DIABLO-selected Component 1 features across models; size indicates absolute loading value, color indicates direction, and shape indicates omics layer.

### Validation in Human Donor Lung Reduced CLOCK Gene Expression in AKI

To test whether our findings apply to human disease, we studied lung tissues from organ donors collected by LifeLink. Donors were divided into AKI and Control groups based on their highest blood creatinine levels measured in hospital (with AKI defined as creatinine≥ 1.3 mg/dL, Table 1). Using RT-qPCR, we examined three biomarkers identified from our multi-omics analysis. While FOXF1 (Figure 9A) and ATF6 (Figure 9B) showed no significant changes, we observed a significant decrease in CLOCK (Figure 9C) gene expression in lungs from AKI donors. Overall, these human data confirm that AKI rapidly disrupts the lung’s circadian regulation, supporting our cross-species findings of altered pulmonary gene activity during AKI-related organ injury.

**Figure 9.**
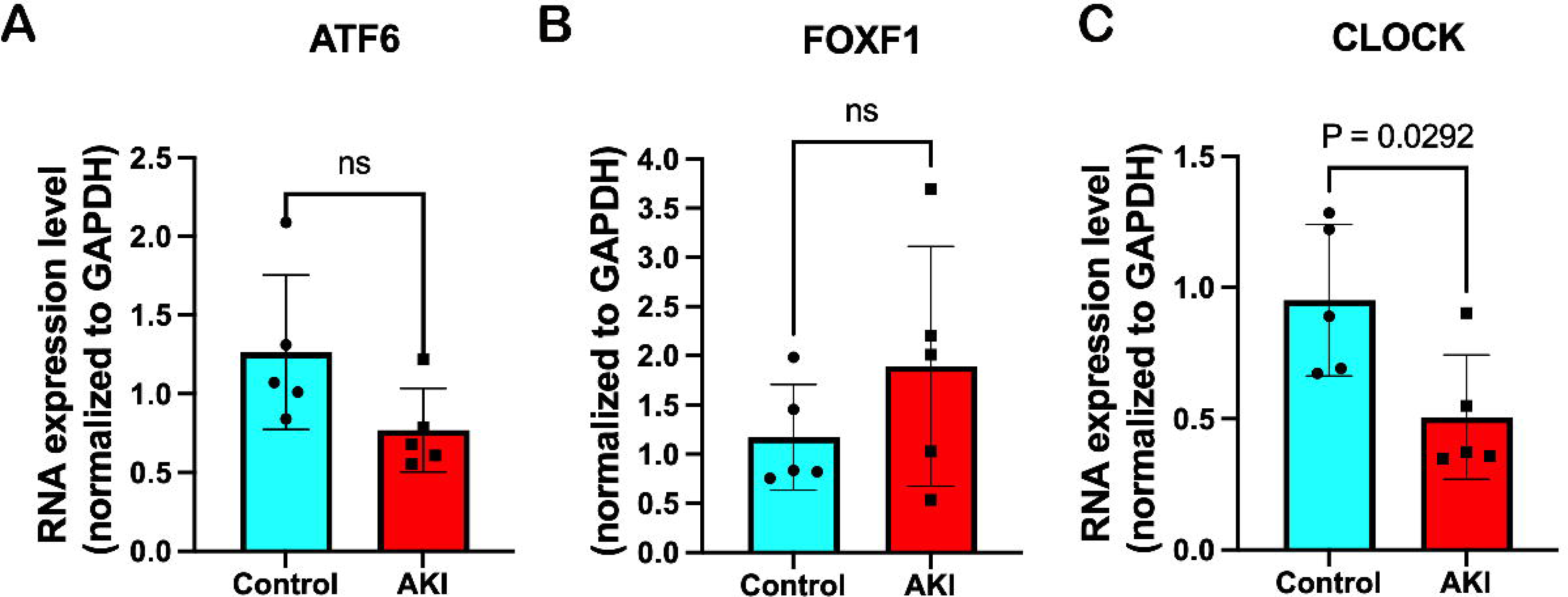
Real time qPCR verification of biomarkers identified by cross species multi omics analysis in primary human donor lung tissue (n=5 in each group). RNA expression levels of (**A**) FOXF1, (**B**) ATF6, (**C**) CLOCK in human lung tissue. Data are expressed as mean ± SD. All data are analyzed using GraphPrism with Student’s unpaired t-test.

**Table 1.**
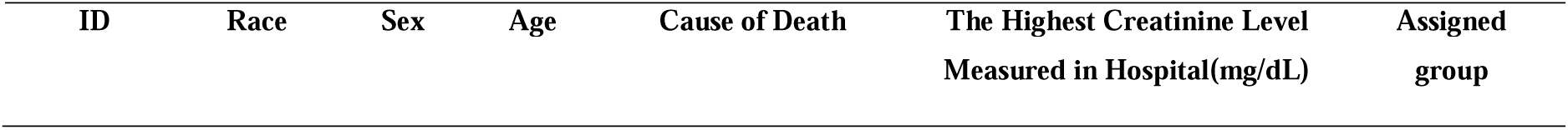
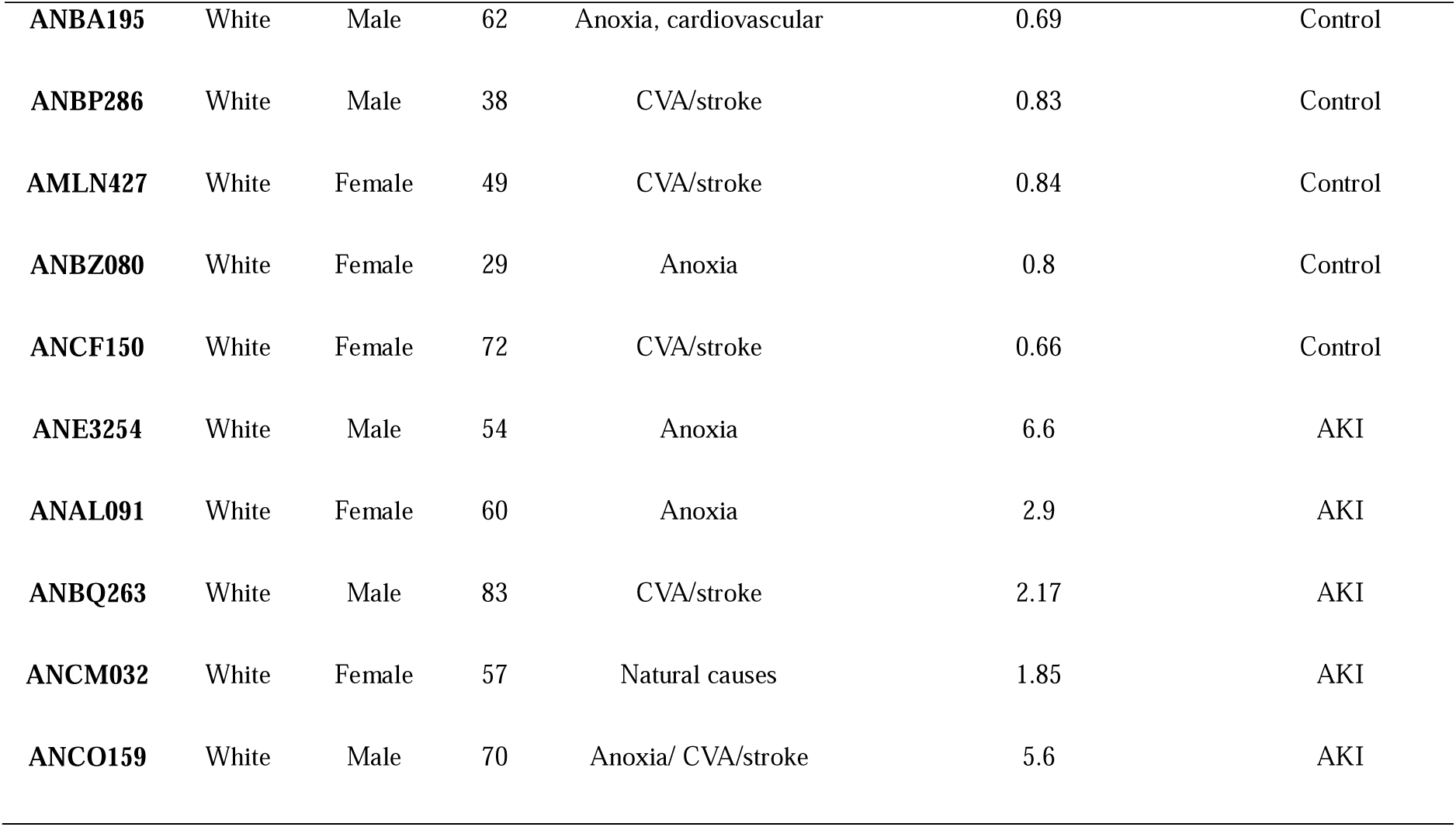
Demographics and Clinical Characteristics of LifeLink Human Lung Donors.

| ID | Race | Sex | Age | Cause of Death | The Highest Creatinine Level Measured in Hospital(mg/dL) | Assigned group |
| --- | --- | --- | --- | --- | --- | --- |
| **ANBA195** | White | Male | 62 | Anoxia, cardiovascular | 0.69 | Control |
| **ANBP286** | White | Male | 38 | CVA/stroke | 0.83 | Control |
| **AMLN427** | White | Female | 49 | CVA/stroke | 0.84 | Control |
| **ANBZ080** | White | Female | 29 | Anoxia | 0.8 | Control |
| **ANCF150** | White | Female | 72 | CVA/stroke | 0.66 | Control |
| **ANE3254** | White | Male | 54 | Anoxia | 6.6 | AKI |
| **ANAL091** | White | Female | 60 | Anoxia | 2.9 | AKI |
| **ANBQ263** | White | Male | 83 | CVA/stroke | 2.17 | AKI |
| **ANCM032** | White | Female | 57 | Natural causes | 1.85 | AKI |
| **ANCO159** | White | Male | 70 | Anoxia/ CVA/stroke | 5.6 | AKI |

## DISCUSSION

In this study, we provide a comprehensive, cross-species multi-omics analysis of AKI-induced lung injury, integrating clinical data with porcine, murine, and human organoid models to map the temporal and molecular landscape of kidney-lung crosstalk. Our findings address a critical knowledge gap in critical care medicine: the sequence by which acute renal injury leads to remote pulmonary dysfunction, progressing from early microvascular and structural changes to transcriptionally mediated inflammation and epithelial stress responses.

Previous studies have reported high mortality in selected intensive care unit (ICU) cohorts ^21,22^. In the present analysis, recorded all-cause mortality was 36.67% in the subgroup with lung injury following AKI and 25.45% in the overall AKI cohort. The reported LOS summaries were also higher in the AKI-lung query than in the AKI-alone query. These observations are consistent with prior evidence linking AKI and pulmonary complications to adverse outcomes ^10,23–25^, and provide clinical context for the experimental findings. However, they do not demonstrate that lung injury independently increases mortality or LOS.

Bioinformatics approaches have revolutionized modern biomedicine, offering unprecedented power to decipher complex disease mechanisms, resolve cellular heterogeneity, and map complicated signaling networks across organ systems ^26–28^. However, applying these computational pipelines to a single layer of biological data will cause bias that can diminish primary pathological drivers. Transcriptomic profiling alone operates on the assumption that mRNA abundance predicts protein expression. Conversely, relying exclusively on proteomics can miss downstream transcriptional adaptation, epigenetic regulation, and long-term gene-regulatory networks. To overcome these single-layer limitations, integrative transcriptomic and proteomic approaches have emerged as powerful strategies for capturing complementary layers of gene regulation along the DNA–RNA–protein axis ^29–32^. Our findings strongly validate this dual-layer strategy. In our porcine model of hyper-acute AKI-induced lung injury, the transcriptomic profile remained stable, exhibiting minimal functional gene expression changes during early reperfusion. If our bioinformatics analyses relied only on RNA-seq, this hyper-acute phase would have been falsely identified as transcriptionally inactive. Instead, parallel proteomic profiling revealed extensive pre-transcriptional remodeling characterized by rapid ciliary, cytoskeletal, and metabolic protein degradation, and a near-zero RNA-protein correlation (r = −0.06). By combining both modalities, we eliminated single-omics bias and demonstrated that early organ crosstalk is driven by immediate post-translational events long before classic transcriptional programs are engaged.

To discovery the universal drivers of AKI-induced lung injury while overcoming the physiological limitations of any single experimental model, we deployed a cross-species approach integrating porcine, murine, and human organoid systems. Experimental models of inter-organ communication are essential for mechanistic investigation, yet each possesses translational limitations. Rodents are the most widely used model for analyzing immune and molecular pathways, but they are different from human in pulmonary anatomy, respiratory mechanics, and immune responses, which limit their capacity to fully model human lung pathophysiology^33–35^. Conversely, porcine models provide a more accurate representation of human cardiovascular and pulmonary physiology, making them particularly useful for investigating hyperacute vascular responses relevant to critical illness. However, compared with murine models, their experimental utility remains restricted by more limited feasibility of genetic engineering and fewer well-established molecular tools^36–39^. Finally, human lung organoids overcome many of the species-specific limitations of animal models and enable direct investigation of airway epithelial stress responses. However, they lack a functional vasculature, circulating immune cells, and the complex hemodynamic and inter-organ interactions that characterize human critical illness ^40,41^. By bridging these three platforms, our framework combined the physiological fidelity of the pig to capture early post-translational barrier failure, the genetic and temporal depth of the mouse to resolve downstream transcriptional inflammation, and human organoids to validate localized epithelial stress axes without systemic confounders. This cross-species strategy provided a highly rigorous and translationally relevant roadmap for therapeutic targeting in human patients.

A central finding of this study is that AKI-induced lung injury progresses through distinct regulatory phases, characterized by a fundamental shift in central dogma coupling over time. In our 4-hour porcine model, transcriptomic changes were virtually undetectable by PCA, showing a muted mRNA profile with only 30 DEGs. However, parallel proteomic profiling revealed a distinct, significant separation driven by 112 DEPs and a statistically significant negative global correlation. This temporal decoupling highlights that the initial, hyper-acute phase of distant lung injury is governed primarily by post-translational mechanisms, proteasomal degradation, and the mobilization or consumption of pre-existing protein pools rather than de novo transcription. Biologically, this early phase is dominated by targeted structural/ciliary protein loss and metabolic reprogramming.

As the injury progresses to the established phase, captured by our 24-hour murine model, the molecular machinery shifts toward a highly coordinated transcriptional program. This transition is marked by a broad transcriptomic shift (2,384 DEGs) and a significant positive proteogenomic correlation (r = 0.316; increasing to r = 0.447 across shared proteins) reflecting efficient translation of inflammatory effectors. Here, active recruitment of classical inflammatory pathways occurs alongside a simultaneous failure of essential homeostatic.

To bridge these animal models directly to human biology without multi-organ systemic confounders, we treated NHBE-derived human lung organoids with conditioned media from mtDAMP-stimulated macrophages. At 24 hours post-treatment, the human organoids exhibited low RNA-protein concordance (r = 0.035) alongside widespread transcriptional suppression of ECM organization, cell motility, and FGF signaling. This transcriptional shutdown indicates that by 24 hours, the human airway epithelium enters a distinct injury-resolution and tissue-repair phase in response to indirect, macrophage-mediated DAMP signaling. Notably, while structural remodeling genes were suppressed, upregulated genes were exclusively enriched for ER stress and UPR pathways. This indicates that while the immediate tissue-damaging signal decays, persistent proteostatic and ER stress remains a long-term cellular liability in human airway cells exposed to mtDAMP.

To extract meaningful biological signatures from complex, multi-layered data across three distinct species, we combined an advanced bioinformatics tools including MOFA+, decoupleR (incorporating PROGENy and CollecTRI), and DIABLO. Our use of unsupervised multi-omics integration through MOFA+ enabled the identification of concealed sources of variation shared across transcriptomic and proteomic datasets without imposing prior biological assumptions ^42–44^. Despite substantial differences in species, injury duration, and experimental models, this approach revealed a conserved latent factor, which highlighting a common molecular response to AKI-induced lung injury. We have isolated clear model-specific signals, capturing mouse-specific inflammation (Factor 1), human organoid ER responses (Factor 2), and pig structural stress (Factor 3). Factor 6 was heavily enriched for vascular remodeling and epithelial repair programs, while Factor 7 captured persistent blood coagulation and hemostatic dysregulation, establishing microvascular remodeling and hemostatic dysfunction as core, species-independent features of AKI-lung crosstalk.

To move beyond latent statistical factors toward mechanistic interpretation, we applied decoupleR to infer upstream signaling pathway activities (PROGENy) and transcription factor regulatory networks (CollecTRI) directly from transcriptomic data. This activity-based framework enabled the identification of upstream regulatory programs that were not readily apparent from differential gene expression alone, particularly in models where transcriptional changes were modest or influenced by post-transcriptional regulation ^45–48^. PROGENy pathway analysis confirmed a progression from silent/subtle transcriptomic pathway activity at 4 hours in pigs to robust activation of NFκB, TNF-α and hypoxia at 24 hours in mice. Using decoupleR with the CollecTRI database, we identified key upstream transcription factors across models. While pig early stress was driven by mesenchymal/structural TFs (FOXF1, PRRX1) and human organoids were driven by UPR TFs (ATF6, ATF2, ATF4), FOXF1, a master regulator of pulmonary endothelial and mesenchymal integrity, was conserved across all three systems, reinforcing the vascular niche remodeling signature highlighted by MOFA+ Factor 6. Furthermore, the conserved activation of SMAD3 and SMAD7 across models pinpoints TGF-β signaling modulation as a fundamental feature of the AKI-lung axis.

DIABLO provided a complementary supervised framework for prioritizing correlated cross-omics biomarkers associated with distinct injury stages across species ^30,49^. Supervised multi-block discriminant analysis using DIABLO yielded high classification accuracy across models (80% in pig, 100% in mouse). Importantly, no biomarkers were shared across all three models, indicating that AKI-induced lung injury does not possess a single static biomarker signature. Instead, DIABLO confirmed that each model captures a distinct, phase-specific molecular state along the continuum of organ injury, from early ciliary/metabolic protein loss (IFT25, PDHA1) to established fibrotic/inflammatory drivers (LOX, BPIFA2) and resolution-phase ECM suppression (FAP, POSTN).

The complementary use of unsupervised latent factor modeling (MOFA+), regulatory activity inference (decoupleR), and supervised multi-block integration (DIABLO) enabled us to move from data-driven discovery to mechanistic interpretation and biomarker prioritization. Together, these orthogonal analytical frameworks increased confidence that the identified molecular signatures represent conserved biological responses to AKI-induced lung injury rather than artifacts of individual datasets or experimental models.

A key result of our study is the marked decrease in CLOCK gene expression, a central circadian regulator, in lung tissue from human donors with AKI. Although kidney failure can affect the whole body, it is less clear how AKI disrupts gene activity in distant organs like the lungs. The CLOCK gene is important for maintaining the lung’s internal rhythms, which helps control blood vessel stability and immune cell movement. When this rhythm is disrupted, the lung’s blood vessels become more fragile and prone to inflammation ^50,51^. Our findings in human tissue directly link the rapid molecular breakdown seen in our pig model to real clinical AKI. In pigs, acute kidney injury led to a rapid loss of vascular barrier integrity and metabolic changes. The parallel reduction of CLOCK in both humans and pigs suggests that signals from the injured kidney, such as mitochondrial DAMPs and inflammatory factors, can actively disrupt the lung’s own biological clock. This may weaken the lung’s ability to repair itself and make it more vulnerable to further damage.

Although our human sample size was small, the significant drop in CLOCK expression among AKI donors points to a strong biological effect. Combined with our analysis of more than 13 million AKI patients in the Epic Cosmos cohort, these results show that AKI-induced lung injury is an active process involving changes in local circadian and metabolic control, not just a passive result of fluid overload. Restoring circadian signaling in the lung could be a promising new approach to protect against organ damage after AKI. Our work also highlights the value of using both transcriptomics and proteomics together. If we had used only gene expression data, we would have missed the early changes seen in the pig model. If we had used only protein data, we would not have detected the later gene-level shutdown and stress responses in the human organoid model. By comparing across species and data types, we captured the full sequence of injury and repair, bridging the gap between experimental models and human disease.

## CONCLUSION

In summary, our study shows that lung injury caused by AKI is a major factor in patient deaths and longer hospital stays. By analyzing data from 13.8 million AKI patients and combining this with detailed molecular studies in animal models and human tissues, we found that AKI-related lung damage develops in distinct stages. Early on, injury is driven by rapid changes in blood vessel function and protein-level metabolism. Importantly, our analysis of human donor lungs confirmed that AKI leads to a clear decrease in the CLOCK gene, which controls the lung’s natural daily rhythms. This effect is directly related to how severe the kidney injury is and does not depend on factors like age, sex, race. These results change the view of AKI-induced lung injury: rather than being just a result of fluid overload, it is an active process involving blood vessel failure and changes in the lung’s internal clock. Identifying these shared molecular changes provides new, human-validated targets to help prevent or treat lung injury in patients with AKI.

## DATA AVAILABILITY STATEMENT

All raw and processed RNA-seq and proteomics datasets generated in this study have been deposited in public repositories. Sequencing data are accessible via the Gene Expression Omnibus (GEO) under accession GSEXXXXXX, and mass spectrometry proteomics data are deposited to the ProteomeXchange Consortium via the PRIDE partner repository under dataset identifier PXDXXXXXX [datasets will be made fully public upon peer-reviewed publication

## ACKNOWLEDGMENTS

This work was supported, in part, by the National Institute of Diabetes and Digestive and Kidney Disease of the National Institutes of Health under award numbers R01DK138092, R01DK134000, and R01DK134028.

The content is solely the responsibility of the authors and does not necessarily represent the official views of the National Institutes of Health.

## AUTHOR CONTRIBUTIONS

Z.T. and R.L. designed experiments presented in the manuscript. X.S. performed the clinic outcomes analysis. Z.T., N. S., D.A-G., G. A. N.Z., L.L., D. R. C., N.H., K. W, R. N., Y. T., S. C., V. S., K.D., L.L., I. L., C. E, C. P. M. T., L. W., performed the pig IR-AKI surgeries, Z.T. performed the mice IR-AKI surgeries, Z.T., N.S., D.A-G., G.A., and Y.T. performed the human lung organoids experiments. Z.T. analyzed the data presented in the manuscript. Z.T. wrote the manuscript. N. S., D.A-G., G. A. N.Z., D. R. C., N.H., K. W, R. N., V.S., K. D., L.L., and R.L. reviewed and edited the manuscript. All authors read and approved this manuscript for submission.

